# Active site structure of the *Shigella flexneri* effector OspI

**DOI:** 10.1101/2022.02.15.480433

**Authors:** Akira Nishide, Kenji Takagi, Minsoo Kim, Tsunehiro Mizushima

## Abstract

Ubc13 is a critical ubiquitin-conjugating enzyme involved in the nuclear factor-κB (NF-κB) signalling pathway. The *Shigella flexneri* effector OspI targets the host Ubc13 and modifies this enzyme by deamidation of Gln100 into Glu100. This modification inhibits the tumour necrosis factor (TNF) receptor-associated factor 6 (TRAF6)-catalyzed ubiquitination and diacylglycerol-CBM (CARD–Bcl10– Malt1)-TRAF6-NF-κB signal activation. We have previously reported the wild-type OspI crystal structure, but the catalytic triad does not form the canonical active site. Here, the crystal structure of OspI with a C62S mutation was determined at a resolution of 2.2 Å. This C62S mutant structure provided the active site conformation with the catalytic site of OspI.

## 1. Introduction

Protein ubiquitination is a critical mechanism in regulating numerous eukaryotic cellular processes, including cell cycle progression, transcriptional control, protein trafficking, signal transduction, immune responses, cancer and infectious diseases. Ubiquitination is catalyzed by a sophisticated cascade system consisting of ubiquitin-activating (E1), ubiquitin-conjugating (E2) and ubiquitin-ligating (E3) enzymes. The E1 enzyme forms a thioester linkage between its active site cysteine residue and ubiquitin. Ubiquitin is next transferred to a cysteine residue on the E2 enzyme. The E2 enzyme binds one of several E3 enzymes, and the E3 enzyme transfers ubiquitin from E2-ubiquitin to the lysine residue of a substrate protein (Weissman *et al.*, 2011). E2 enzymes play a central role in the enzymatic cascade of ubiquitination. Ubc13 is unique among E2 enzymes and forms a heterodimeric complex with a ubiquitin E2 variant (Uev1A). Ubc13 and Uev1A are responsible for the non-canonical K63-linked ubiquitination of tumour necrosis factor (TNF) receptor-associated factor (TRAF)-family adapter proteins involved in Toll-like receptor and TNF-family cytokine receptor signalling, which are regulators of innate immunity. Particularly, Ubc13-induced TRAF6 ubiquitination is linked to the activation of the NF-κB signalling pathway, resulting in the activation of immune and inflammatory responses and protection against apoptosis (Chen, 2005).

Many pathogenic bacteria deliver virulence factors, called effectors, into host cells through the type III secretion system. Recent studies have identified several bacterial effectors that interact and modulate the ubiquitination pathway during pathogenic bacteria infection. Bacterial effectors mimic enzymes of the host ubiquitin system or bind and modify components of the host ubiquitin system and modulate host signalling pathways to promote bacterial infection (Kim *et al.*, 2010; Kim *et al.*, 2014). OspI, a *Shigella flexneri* effector, specifically binds to Ubc13 and modifies this enzyme by deamidation of Gln100 into Glu100 (Sanada *et al.*, 2012). This modification inhibits TRAF6-catalyzed ubiquitination and activation of the diacylglycerol-CBM-TRAF6-NFκB signalling pathway (Sanada *et al.*, 2012, Mohanty et al. 2019). We have determined the crystal structures of OspI alone and OspI C62A in complex with Ubc13 (Sanada *et al.*, 2012; Nishide *et al.*, 2013). OspI has a cysteine protease-like fold and three conserved active site catalytic residues (Cys62, His145 and Asp160). However, the Sγ position of the Cys62 in wild-type OspI was located on a different side of the active site in the canonical cysteine protease superfamily (Sanada *et al.*, 2012). Although the active site structure of OspI C62A (Nishide *et al.*, 2013; Fu *et al.*, 2013) has been observed to undergo conformational changes, the precise spatial arrangement of the active site residues remains unknown. To define the active site structure of OspI precisely, we determined the crystal structure of the OspI C62S mutant.

## 2. Materials and methods

### 2.1. Expression, purification and crystallization

*S. flexneri* OspI with the C62S mutation was cloned into the pGEX6P-1 vector and expressed in *Escherichia coli* BL21. The plasmids were used to transform the BL21 strain of *E. coli*, and protein expression was induced by adding 0.1 mM isopropyl-b-D-thiogalactopyranoside (IPTG) at 37°C for 3 h. Cell extracts were prepared by sonicating in lysis buffer (20 mM Tris-HCl, pH 7.4, and 150 mM NaCl) at 27,000 *×g* for 20 min at 4°C. GST fusion proteins were purified by Glutathione Sepharose 4B and gel filtration chromatography (Superdex 75, Cytiva). The GST moiety was removed by cleaving with the PreScission protease and subsequently performing gel filtration chromatography (Superdex 75, Cytiva). The purified proteins were concentrated to 21.9 mg/mL by ultrafiltration in 25 mM Tris-HCl (pH 7.5) and 1 mM dithiothreitol. Crystallization of the OspI C62S mutant was performed using the sitting-drop vapour diffusion method at 293 K in drops containing a mixture of 1μL of protein solution and 1 μL of reservoir solution, which consisted of 0.2 M magnesium acetate, 0.1 M sodium cacodylate (pH 6.5) and 20% PEG8000.

### 2.2. X-ray data collection, structure determination and refinement

The crystals were frozen without cryoprotective additives. X-ray diffraction datasets were collected at 100 K on beamline BL44XU at SPring-8 (Hyogo, Japan). The C62S mutant crystals diffracted to maximum resolutions of 2.01 Å. Data processing and reduction were performed using HKL-2000 (Otwinowski & Minor, 1997). The crystals belonged to space group *C*2221, with one molecule in the asymmetric unit, corresponding to a Matthews coefficient of 2.52 Å^3^ Da^−1^ (51.2% solvent content) (Matthews, 1968). The crystal structure of the OspI C62S mutant was determined by molecular replacement in MOLREP (The CCP4 suite: programs for protein crystallography, 1994, Vagin & Teplyakov, 2010) using the wild-type OspI (PDB ID code 3B21) structure lacking residues 59–77 as the search model. The models were subsequently improved through alternate cycles of manual rebuilding using COOT (Emsley & Cowtan, 2004) and refinement with the program REFMAC5 (Murshudov *et al.*, 1997). Water molecules were built by ARP/wARP (Morris *et al.*, 2003). Model validation was performed using PROCHECK (Laskowski *et al.*, 1993) from the CCP4 suite. There were no residues in disallowed regions of the Ramachandran plot. The structure figures were generated using CCP4MG (McNicholas *et al.*, 2011). Data processing and refinement statics are summarised in Table 1.

**Table 1.** Data collection and refinement statistics

|  |  |
| --- | --- |
| **Data collection** |  |
| Space group | $C222_1$ |
| Unit-cell parameters |  |
| $a$ (Å) | 68.4 |
| $b$ (Å) | 92.0 |
| $c$ (Å) | 68.1 |
| Wavelength (Å) | 0.9 |
| Distance (mm) | 250 |
| Exposure time (sec.) | 1 |
| Resolution range (Å) | 50.0–2.20 (2.24–2.20) |
| No. of total reflections | 80260 |
| No. of unique reflections | 11184 |
| Completeness (%) | 99.9 (100.0) |
| $R_{\text{merge}}$ (%)† | 9.0 (24.5) |
| $I/\sigma(I)$ | 14.3 |
| Redundancy | 7.2 (7.4) |
| **Refinement** |  |
| Resolution (Å) | 38.11–2.20 (2.26–2.20) |
| No. of reflections | 10572 |
| $R_{\text{cryst}}$ (%) | 19.0 (23.6) |
| $R_{\text{free}}$ (%) | 25.2 (34.7) |
| R.m.s.d. bond lengths (Å) | 0.010 |
| R.m.s.d. bond angles (°) | 1.251 |
| No. of protein atoms | 1491 |
| No. of water | 110 |
| Average $B$ factors (Å<sup>2</sup>) | |
| Protein | 10.7 |
| Solvents | 15.7 |
| Ramachandran plot (%) |  |
| Most favored region | 90.6 |
| Additional allowed | 9.4 |
| Generously allowed | 0.0 |

† $R_{\text{merge}} = \sum_h \sum_i |I_{hi} - \langle I_h \rangle| / \sum_h \sum_i I_{hi}$ , Values in parentheses are for the outer shell.

Structure factors and coordinates have been deposited in the Protein Data Bank (PDB; http://www.rcsb.org/pdb) under the accession code 4XZX.

## 3. Results and Discussion

### 3.1. Overall structure of the OspI C62S mutant

The crystal structure of the OspI C62S mutant was refined at a resolution of 2.2 Å. The final refined model comprises residues 21–212 of an OspI C62S molecule in the asymmetric unit. The N-terminal 20 residues (1–20) are missing in the final model because of the lack of electron density. The structure of the OspI C62S mutant, which consists of seven α-helices (α1–α7), five β-strands (β0–β4) and a 310 helix (310-1), is essentially identical to wild-type OspI (PDB ID code 3B21) (Fig. 1a). However, the α4–α5 loop region forms an extra short β-strand (β0) leading to the formation of five stranded β-sheets with β1–β4. The superposition of these structures showed that the largest shifts of Cα atoms are located in the N-terminus of α3, with and overall root mean square deviation (r.m.s.d.) of 0.20 Å on Cα atoms (Fig. 1b, c). The active site Cys62 in OspI is located at the N-terminus of α3. This result suggests that the active site conformation is changeable in OspI.

**Figure 1.**
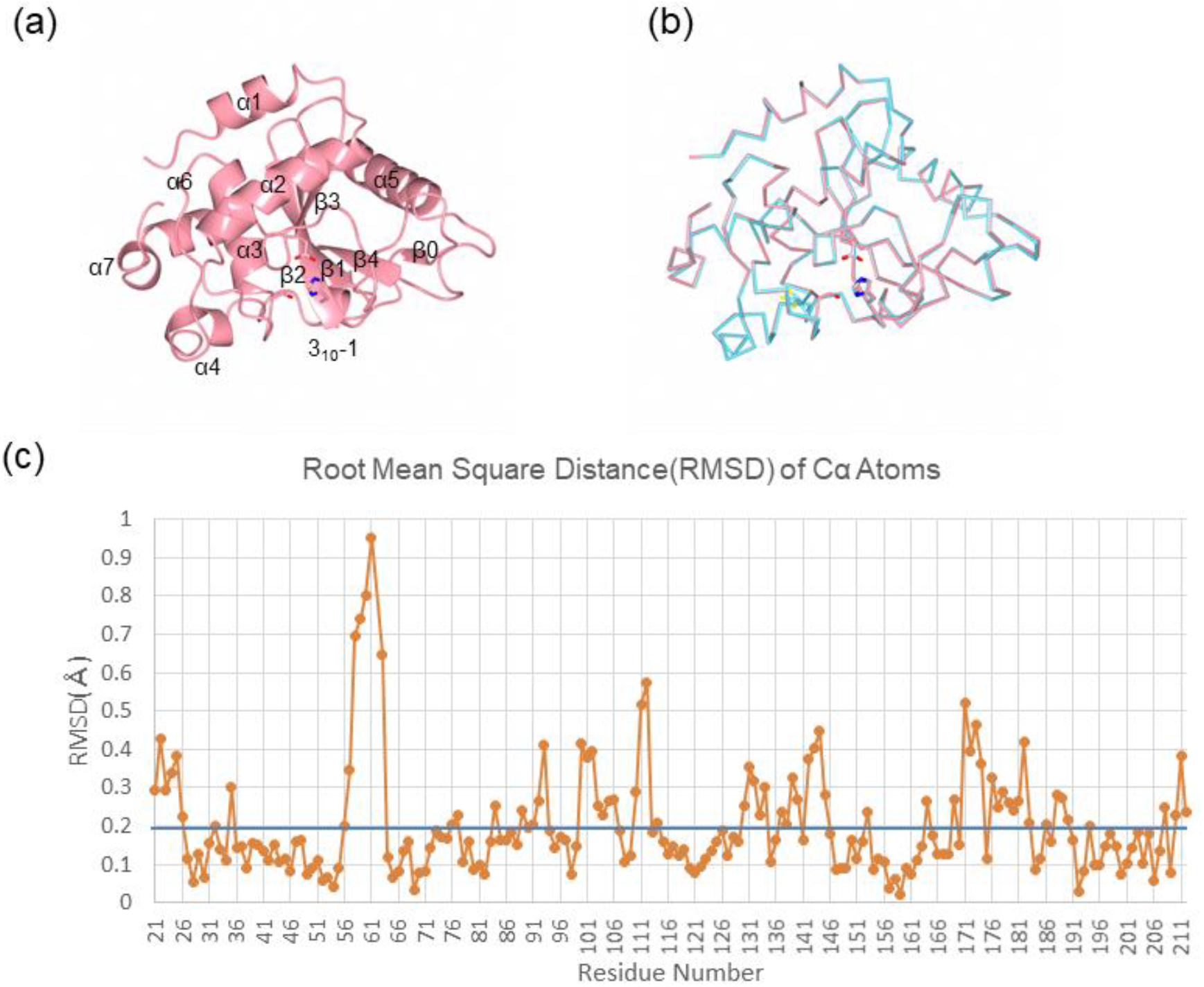
Structure of the *Shigella flexneri* OspI C62S mutant. (a) Ribbon diagram of the OspI C62S mutant. The active site residues are shown as stick models. (b) Comparison between the wild-type (cyan; PDB ID code 3B21) and the C62S (pink; PDB ID code 4XZX) OspI structures. Molecules were superimposed using the Cα positions. (c) R.m.s.d. on Cα atoms by residues between the OspI C62S and wild-type OspI structures

### 3.2. Structure of the active site

A cysteine protease-like catalytic triad was identified in the structure of OspI and comprises residues Cys62, His145 and Asp160. This triad is essential for the deamidase activity of OspI, and a mutation in any of these residues leads to the loss of its activity. However, Cys62 exists in three discrete conformations in the crystal structure, and the highest occupancy site of Cys62 appears to form a disulphide bond with Cys65 (Sanada *et al.*, 2012). Cys62 in these conformers is too far to form a hydrogen bond with His145. The crystal structure of wild-type OspI has an unusual structural element orienting the nucleophilic cysteine.

The structure of the active site triad of the OspI C62S mutant is shown in Fig. 2a. The structure demonstrates that the presence of non-protein electron density map at the active site, which is modelled as acetate according to the shape of the electron density and crystallization conditions (Fig. 2a). The active site cysteine is located at the N-terminus of the α3 helix with the side chains of His145 and Asp160 arising from the β2 and β3 strands, respectively. In the OspI C62S mutant, the residues of the catalytic triad are connected through hydrogen bond interactions. The hydroxyl oxygen of the catalytic serine (from sulphur in cysteine to oxygen in serine) is positioned at 3.2 Å from the Nδ1 atom of His145. The orientation of His145 is stabilised by a hydrogen bond from the Nε2 atom of this residue to the Oδ1 atom of Asp160 (2.7Å) (Fig. 2a). These inter-residue distances are similar to those found in cysteine protease AvrPphB (Zhu *et al.*, 2004). This spatial arrangement will presumably enhance the nucleophilicity of the active site cysteine in OspI in a similar manner to that observed in this archetypal superfamily of enzymes including cysteine proteases, acetyl transferases, deamidases, transglutaminases and ubiquitin carboxyl-terminal hydrolase (Fig. 2a).

**Figure 2.**
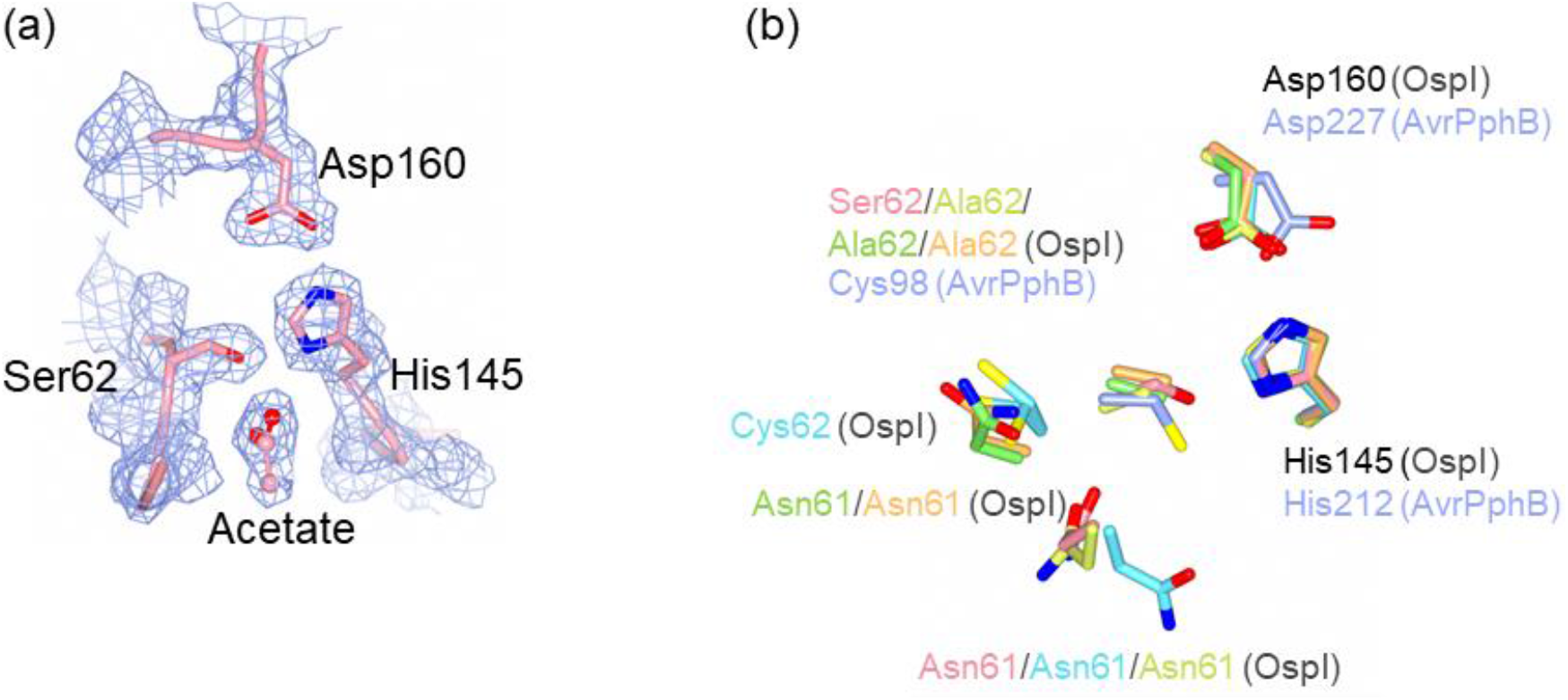
Catalytic active site structure of OspI. (a) 2Fo–Fc electron density map of the active site contoured at 0.5 σ with the final model of OspI C62S superimposed. (b) Comparison of active sites of OspI C62S (pink; PDB ID code 4XZX), wild-type OspI (cyan; PDB ID code 3B21), OspI C62A (yellow; PDB ID code 3W30), OspI C62A bound to Ubc13 (green; PDB ID code 3W31), OspI C62A bound to Ubc13 (orange; PDB ID code 4IP3) and AvrPphB (light blue; PDB ID code 1UKF). Histidines of the catalytic triad were superimposed.

We next compared the structures of the OspI C62S mutant with the known structures of the wild-type OspI and OspI C62A mutants. Although the overall structure of OspI was almost identical among these mutants (r.m.s.d. range for 21–212 Cα atoms of 0.20–0.92 Å), the structures of the active site were classified into three types: wild-type OspI, OspI Cys62 mutants (C62S and C62A) alone and OspI C62A–Ubc13 complex. Compared to the wild-type alone structure, the OspI mutants undergo marked conformational changes. Although the catalytic pocket in wild-type OspI is shielded by Asn61, Asn61 in OspI mutants alone and its complex exhibits an open conformation of the active site (Fig. 2b). In OspI C62S, the side chain of Asn61 forms a hydrogen bond with Ser63. After these conformational changes, the active site triad could be well superimposed with that of AvrPphB (Fig. 2b).

In summary, we determined the crystal structure of the OspI C62S mutant. The refined structure demonstrated that the catalytic pocket of OspI could have different conformations. This structural flexibility may affect the function of OspI.

## Acknowledgements

We thank Chihiro Sasakawa (Chiba University) for helpful discussions. This study was performed using a synchrotron beamline BL44XU at SPring-8 under the Cooperative Research Program of the Institute for Protein Research, Osaka University (proposal 2011A6645, 2011B6645 and 2012A6744). This work was supported by JSPS-KAKENHI JP20H03198 (T.M.), 20H02878 (M.K.), and 19K15759 (A.N) and JSPS Research Fellowships for Young Scientists [14J10879 (A.N.)].

## References

The CCP4 suite: programs for protein crystallography (1994). 50, 760–763.

Chen, Z. J. (2005). Nat Cell Biol 7, 758–765.

Emsley, P. & Cowtan, K. (2004). Acta Crystallogr D Biol Crystallogr 60, 2126–2132.

Fu, P., Zhang, X., Jin, M., Xu, L., Wang, C., Xia, Z. & Zhu, Y. (2013). PLoS Pathog 9, e1003322.

Kim, M., Ashida, H., Ogawa, M., Yoshikawa, Y., Mimuro, H. & Sasakawa, C. (2010). Cell Host Microbe 8, 20–35.

Kim, M., Otsubo, R., Morikawa, H., Nishide, A., Takagi, K., Sasakawa, C. & Mizushima, T. (2014). Cells 3, 848–864.

Laskowski, R. A., MacArthur, M. W., Moss, D. S. & Thornton, J. M. (1993). J. Appl. Cryst. 26, 283–291.

Matthews, P. C. (1968). Int. J. Soc. Psychiatry 14, 125–133.

McNicholas, S., Potterton, E., Wilson, K. S. & Noble, M. E. (2011). Acta Crystallogr D Biol Crystallogr 67, 386–394.

Mohanty P, Agrata R, Habibullah BI, G S A, Das R. (2019) Elife 22, e49223.

Morris, R. J., Perrakis, A. & Lamzin, V. S. (2003). Methods Enzymol 374, 229–244.

Murshudov, G. N., Vagin, A. A. & Dodson, E. J. (1997). Acta Crystallogr D Biol Crystallogr 53, 240–255.

Nishide, A., Kim, M., Takagi, K., Himeno, A., Sanada, T., Sasakawa, C. & Mizushima, T. (2013). J Mol Biol 425, 2623–2631.

Otwinowski, Z. & Minor, W. (1997). Methods Enzymol. 276, 307–326.

Sanada, T., Kim, M., Mimuro, H., Suzuki, M., Ogawa, M., Oyama, A., Ashida, H., Kobayashi, T., Koyama, T., Nagai, S., Shibata, Y., Gohda, J., Inoue, J., Mizushima, T. & Sasakawa, C. (2012). Nature 483, 623–626.

Vagin, A. & Teplyakov, A. (2010). Acta Crystallogr D Biol Crystallogr 66, 22–25.

Weissman, A. M., Shabek, N. & Ciechanover, A. (2011). Nat Rev Mol Cell Biol 12, 605–620.

Zhu, M., Shao, F., Innes, R. W., Dixon, J. E. & Xu, Z. (2004). Proc Natl Acad Sci U S A 101, 302–307.

